# Volume electron microscopy reveals 3D synaptic nanoarchitecture in postmortem human prefrontal cortex

**DOI:** 10.1101/2024.02.26.582174

**Authors:** Jill R. Glausier, Cedric Bouchet-Marquis, Matthew Maier, Tabitha Banks-Tibbs, Ken Wu, Jiying Ning, Darlene Melchitzky, David A. Lewis, Zachary Freyberg

**Author notes:** Current address: University of Pittsburgh, Department of Psychiatry, Pittsburgh, PA 15213.

## Abstract

Synaptic function is directly reflected in quantifiable ultrastructural features using electron microscopy (EM) approaches. This coupling of synaptic function and ultrastructure suggests that *in vivo* synaptic function can be inferred from EM analysis of *ex vivo* human brain tissue. To investigate this, we employed focused ion beam-scanning electron microscopy (FIB-SEM), a volume EM (VEM) approach, to generate ultrafine-resolution, three-dimensional (3D) micrographic datasets of postmortem human dorsolateral prefrontal cortex (DLPFC), a region with cytoarchitectonic characteristics distinct to human brain. Synaptic, sub-synaptic, and organelle measures were highly consistent with findings from experimental models that are free from antemortem or postmortem effects. Further, 3D neuropil reconstruction revealed a unique, ultrastructurally-complex, spiny dendritic shaft that exhibited features characteristic of heightened synaptic communication, integration, and plasticity. Altogether, our findings provide critical proof-of-concept data demonstrating that *ex vivo* VEM analysis is an effective approach to infer *in vivo* synaptic functioning in human brain.

## INTRODUCTION

Coordinated, adaptable synaptic signaling underlies core brain processes such as thought, emotion, learning, and memory^1^. Loss or dysfunction of synaptic signaling is proposed as the pathophysiological substrate for severe brain disorders including schizophrenia^2,3^ and autism spectrum disorder^4^, which are characterized by impairments to these core brain processes. Thus, investigating synaptic signaling in the human brain is critical to advance the understanding of both normal synaptic function and the nature of synaptic dysfunction in disease.

The morphological substrate of synaptic communication is the synaptic complex^5,6^, comprised in its most elemental form by a presynaptic axonal bouton apposed to a postsynaptic element, such as a dendritic spine. The synaptic complex and its constituent components can be directly visualized in human brain tissue with electron microscopy (EM) approaches. A substantial literature in experimental, non-human model organisms demonstrates that *in vivo* synaptic function is directly reflected in ultrastructural measures obtained *ex vivo* via EM, and this is especially evident in the glutamate synaptic system. For example, presynaptic active zone size reflects the relative level of axonal bouton activation and glutamate release probability^7–11^. Postsynaptic density (PSD) size is strongly correlated with excitatory postsynaptic potential amplitude^7^ and abundance of AMPA receptors^12–16^. Because synaptic communication represents the largest energy-demanding process in the brain^17–26^, relative ATP demand is reflected by the abundance^25,27,28^, size^27,29^, and morphology^30–33^ of mitochondria within synaptic complexes. Moreover, a fundamental feature of synaptic communication is that pre– and postsynaptic elements act together as a functional unit^34,35^, and EM ultrastructural measures also capture this core aspect of synaptic functioning^36^. For example, presynaptic glutamate release probability and presynaptic active zone size correlate with PSD size^7,8^. Likewise, presynaptic mitochondrial abundance is related to PSD size^33,37^. This wealth of data from experimental models demonstrates that *in vivo* synaptic function can be derived from quantitative EM analysis of *ex vivo* preserved brain tissue. However, whether these synaptic function-ultrastructure relationships are present in human brain is unclear.

Human brain tissue is sourced either from biopsies obtained during neurosurgical interventions or from donations obtained postmortem. However, postmortem donations are the exclusive source for brain tissue from individuals unaffected by brain disorders during life and for individuals with neuropsychiatric disorders like schizophrenia, autism spectrum disorder, or Alzheimer’s disease, which are currently not diagnosed or treated via neurosurgery. Because whole brains are typically obtained, postmortem sources also permit analysis of multiple brain regions from a single subject, as well as analysis of regions not typically obtained during neurosurgery. For example, the dorsolateral prefrontal cortex (DLPFC) is a higher-order, multi-modal association area^38,39^ that is uniquely expanded in humans^40,41^, and is considered a key site of synaptic dysfunction in human-specific psychiatric disorders^42–45^. EM analysis of postmortem DLPFC tissue therefore presents a unique opportunity to investigate synaptic and sub-synaptic impairments present in individuals with psychiatric disorders relative to individuals unaffected by brain disorders. Although EM studies of postmortem human DLPFC have been published [for examples see^46–49^], this earlier work utilized a conventional two-dimensional (2D) approach to study single ultrathin (∼60 nm) sections in order to generate estimates of three-dimensional (3D) features. Furthermore, a Z resolution of ∼60 nm may obscure critical ultra– and nanostructures, even when ultrathin sections are studied in series. Finally, serious concerns persist regarding potential confounding effects of postmortem biological processes on synaptic and sub-synaptic ultrastructural features.

To address these prior limitations, the primary goal of this study was to develop a new strategy to determine how well synaptic nanoarchitecture and relationships are preserved in postmortem human brain, within the context of the surrounding neuropil^50^. We performed a 3D quantitative analysis of glutamate synaptic complexes in postmortem human DLPFC using focused ion beam-scanning electron microscopy (FIB-SEM). This volume electron microscopy (VEM) approach works by using a focused ion beam to remove a sub-10 nm slice of the tissue, followed by imaging of the newly-exposed cross-section face via SEM in a sequential manner until the entire tissue block is imaged^51^. By iterating through these steps, a 3D volume of brain tissue is acquired with ultrafine Z resolution. Using this innovative technology, we imaged and densely-reconstructed a 3D volume of DLPFC with essentially no loss of information at a 5nm milling step-size from an individual with no brain disorders present during life. Taking advantage of the rich data within this 3D brain volume, we completed targeted, 3D reconstructions and volumetric analyses of 50 Type 1 axo-spinous glutamatergic synapses. Quantitative analyses revealed that the fundamental relationships between pre– and postsynaptic components identified in non-human experimental systems is clearly evident in postmortem human brain tissue. Furthermore, dense reconstruction of DLPFC neuropil revealed novel structural organization within the human DLPFC, including a spiny dendritic segment that exhibited unique and complex ultrastructural features indicative of heightened synaptic communication. Overall, our findings provide a critical proof-of-concept that *ex vivo* VEM analysis offers a valuable and informative means to infer *in vivo* functioning of individual synaptic complexes in human brain.

## RESULTS

### Qualitative Assessment of VEM-Imaged DLPFC Layer 3 Neuropil Volume

Qualitative evaluation of the 3D VEM dataset of postmortem human DLPFC layer 3 revealed excellent ultrastructural preservation^48^ (Figure 1; Supplemental Videos 1-2). Cellular and organelle plasma membranes were intact, and these structures had a paucity of swelling, deformation, or other signs of autolysis. Mitochondria exhibited ultrastructural features characteristic of preserved integrity, including organized cristae, homogenous matrix, and globular or elongated morphology^30,48,52^ (Figure 2). For example, the presence of a globular mitochondrion in the axon terminal forming a synapse (Figure 2A), and the elongated mitochondrion in an axon terminal lacking a synapse (Figure 2C), is consistent with the association of distinct mitochondrial morphologies with high or low energy states, respectively.

**Figure 1.**
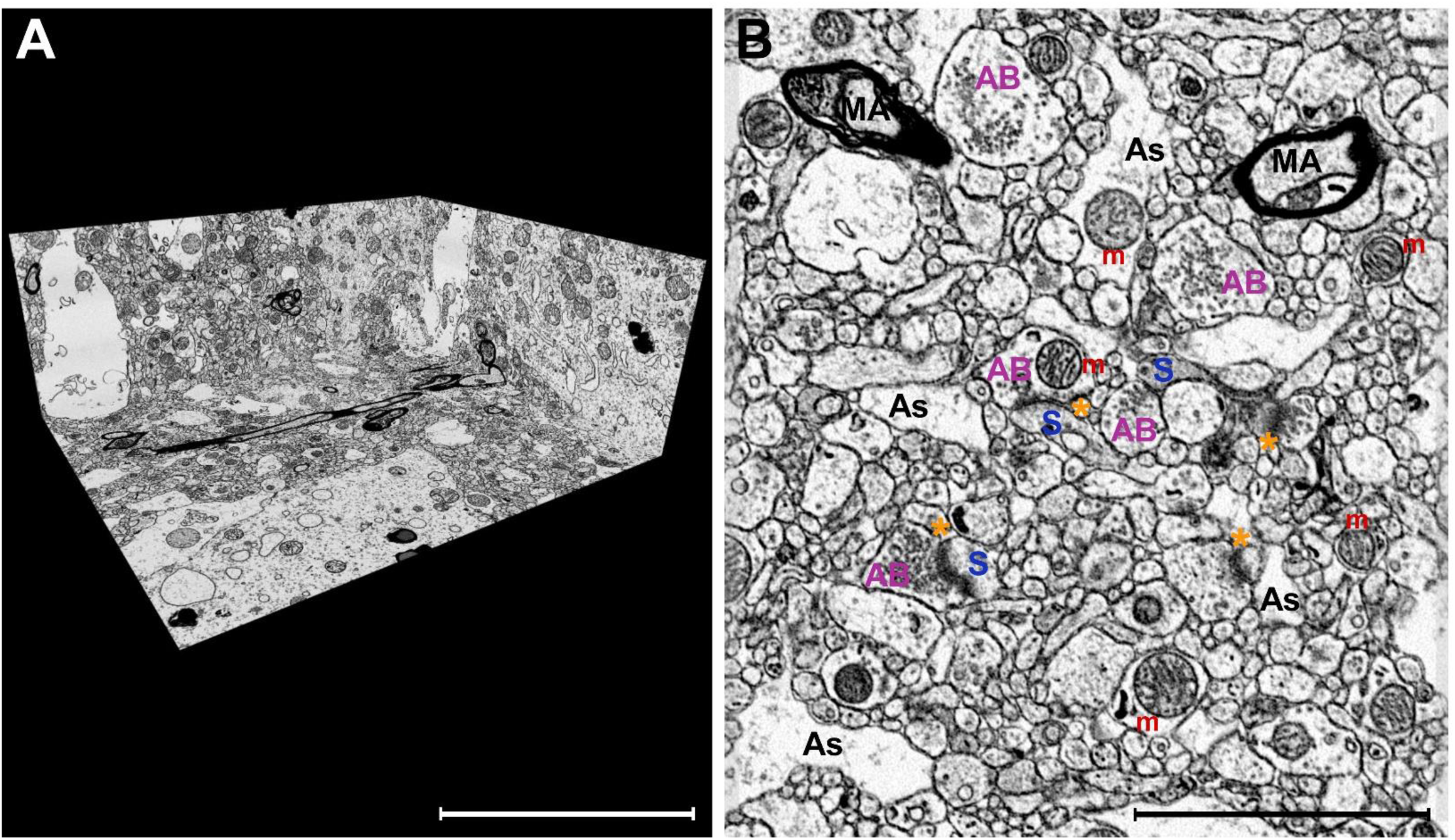
Excellent ultrastructural preservation of postmortem human brain tissue processed for FIB-SEM imaging. **A)** A perspective overview of a volume of DLPFC layer 3 captured with an isotropic voxel size of 5×5×5nm. **B)** A representative SEM image using in-column detector with the Helios 5CX illustrating excellent preservation of the neuropil. Select glutamate synapses labeled: axonal bouton (AB) directly apposed to a dendritic spine (S) containing an electron-dense PSD (asterisk). Additional neuropil components, such as astrocyte processes (As), myelinated axons (MA), and mitochondria (m) are also labeled. Scale bar is 10 μm in A and 3 μm in B.

**Figure 2.**
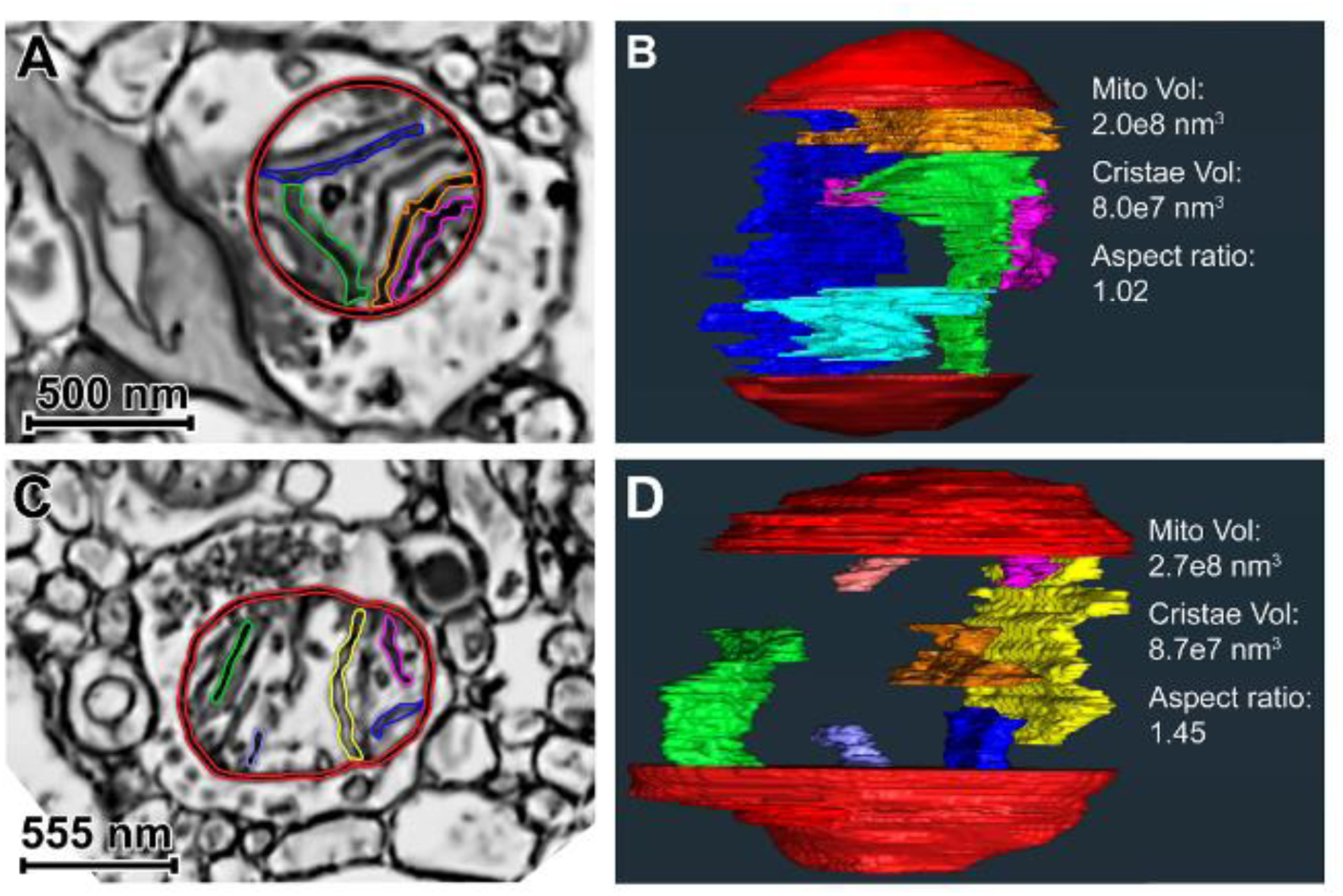
Single SEM images and 3D reconstructions illustrating globular and elongated presynaptic mitochondria in postmortem human DLPFC Layer 3. **A)** Globular mitochondrion (red membrane; multiple-colored cristae). **B)** Internal view of the globular mitochondrion 3D reconstruction, including 5 cristae reconstructions. Total mitochondrion and cristae volumes are provided. The aspect ratio is characteristic of globular morphology. **C)** Elongated mitochondrion (red membrane; multiple-colored cristae) present in a non-synaptic axon terminal. **D)** Internal view of the elongated mitochondrion 3D reconstruction, including 7 cristae reconstructions. Quantitative values of the 3D reconstruction are provided, and the aspect ratio is characteristic of elongated morphology. In Panels B and D, only a subset of cristae volume reconstructions is shown in order to optimize visualization in a 2D image. In Panels A and C, reconstructed cristae visible in the plane of view are colored.

Synaptic complexes and sub-synaptic structures were readily detected in the 3D neuropil volume (Figures 3-4). Dendritic spines were typically the recipient of Type 1 synapses, and these spines often contained a spine apparatus (Figure 3A-B, Figure 4). As expected, mitochondria were not observed in synaptic spine heads, but were observed in the dendritic shaft near the base of the spine neck (Figure 3B), or near Type 1 synapses that formed onto dendritic shafts (Figure 3C). Type 2 synapses, presumably GABAergic, were identified on neuronal somata (Figure 3D). We also observed dually-innervated spines receiving a Type 1 and a Type 2 synapse (Figure 3E-F).

**Figure 3.**
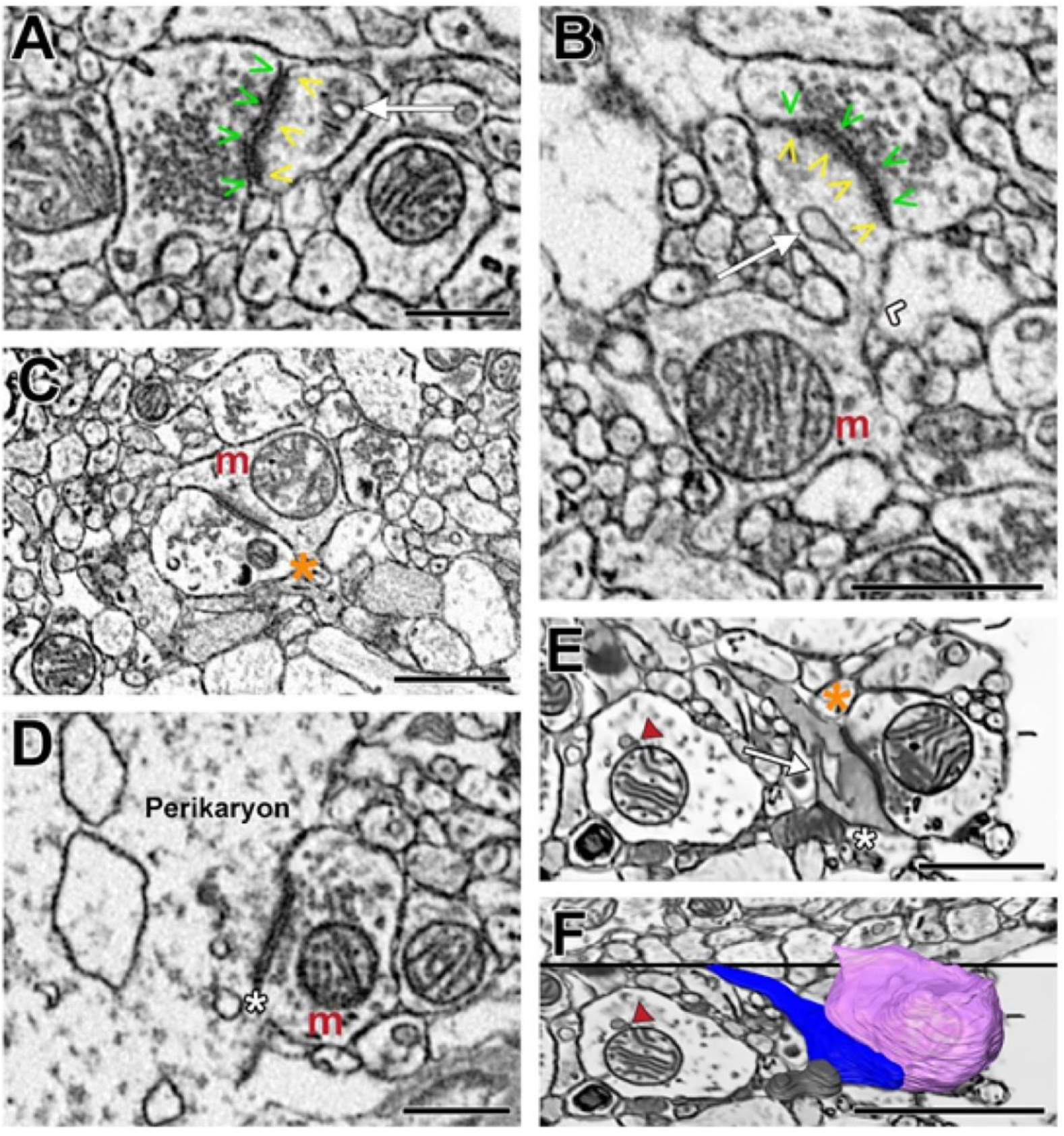
Representative images illustrating Type 1 and Type 2 synapses identified in postmortem human DLPFC layer 3. **A)** Axonal bouton filled with synaptic vesicles and forming a Type 1 synapse onto a dendritic spine containing a spine apparatus. **B)** A parent dendritic shaft containing a mitochondrion positioned at the base of the spine neck. The spine head contains a spine apparatus and is receiving a Type 1 synapse. **C)** A bouton forming a Type 1 synapse onto a dendritic shaft containing a mitochondrion. **D)** Neuronal cell body receiving a Type 2, presumably GABAergic, synapse. **E)** A dendritic spine with a spine apparatus and electron-dense cytoplasm receiving a Type 1 and a Type 2 synapse. In the adjacent dendritic shaft, the mitochondrion is tethered to an electron dense vesicle. **F)** 3D reconstruction of the Type 1 bouton (pink), dendritic spine (blue) and the Type 2 bouton (grey) shown in E within the context of the 2D surrounding neuropil. **Symbol legend:** green arrowheads indicate the active zone; yellow arrowheads indicate the PSD; white arrow indicates the spine apparatus; white chevron indicates the spine neck; orange asterisk indicates a Type 1 synapse; white asterisk indicates a Type 2 synapse; red arrowhead indicates the mitochondrial-tethered vesicle. Scale bar is 1.5μm in A, B, D and F; and is 1μm in E.

**Figure 4.**
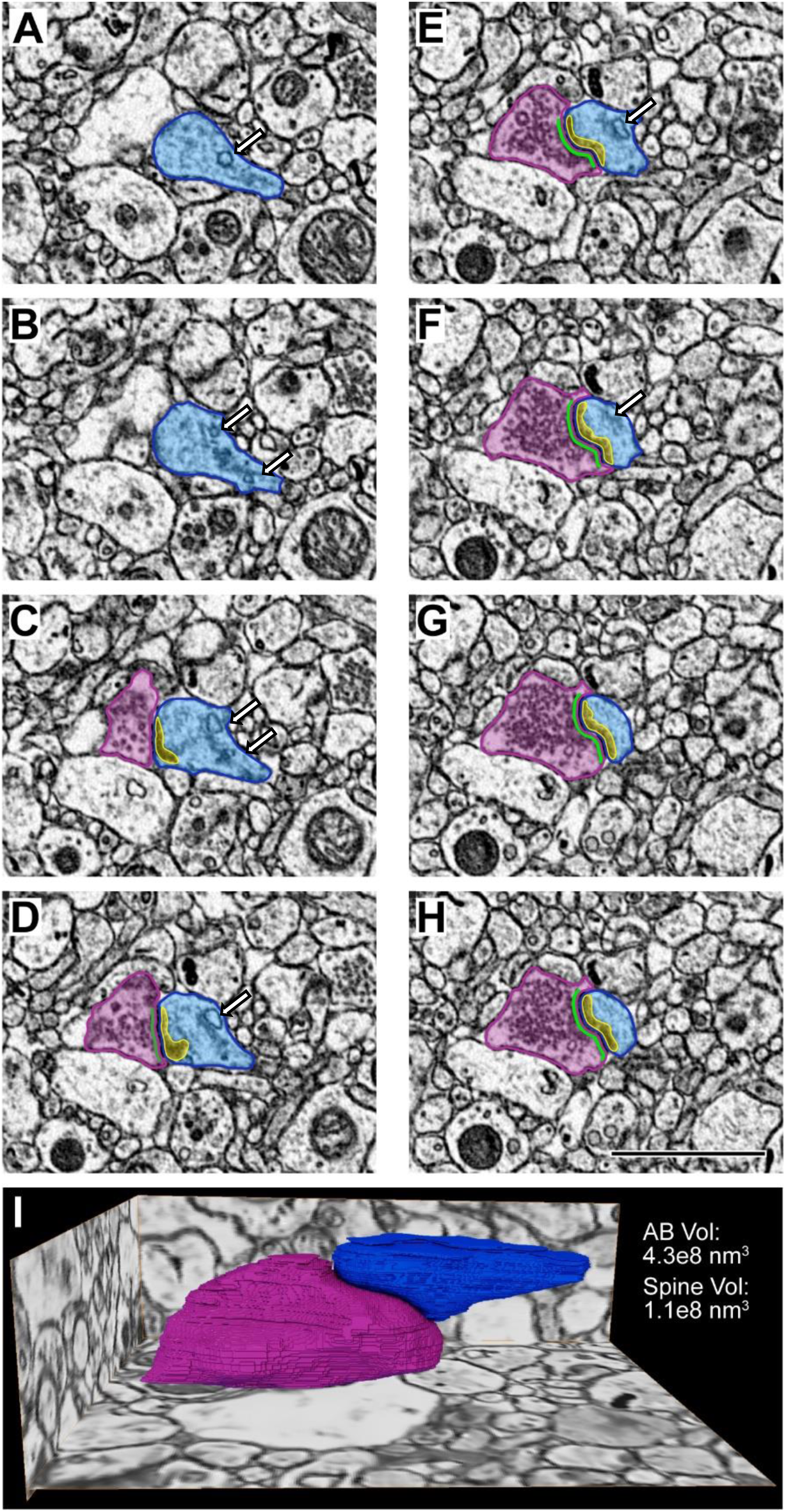
A series of electron micrographs viewed in the XY coordinate plane and the completed volume reconstructions of a Type 1 glutamatergic synapse in postmortem human DLPFC layer 3. **A-H**) Each panel represents 5 nm increments milled by the gallium FIB. In Panels A-B, the dendritic spine (blue) is identified. Panel C is the initial visualization of the glutamatergic synaptic structure including a presynaptic axonal bouton (pink) with an active zone (green) and a postsynaptic spine (blue) containing a PSD (yellow). Note the presence of a spine apparatus (white arrow) in Panels A-F. Scale bar is 2 μm. **I)** Glutamatergic synaptic structure reconstructed in 3D shown in the context of the surrounding tissue. Volume measurements for the axonal bouton (pink) and the spine (blue) are provided.

In qualitatively evaluating the complete 3D volume, we noted a dendritic shaft with distinct sub-cellular structures situated within the cytoplasm, as if piercing through the dendrite. To better interrogate the features and relationships between these structures, we completely reconstructed the shaft in three dimensions, revealing all spines emanating from the shaft, and the sub-cellular structures coursing through the shaft cytoplasm (Figures 5-6). Four distinct, cylindrical structures that were completely engulfed by the dendrite were also reconstructed. Unexpectedly, they did not connect to any parent structure (Figure 5B). On average, these invaginated structures spanned 1.7 ± 0.55 μm in the Z-direction, had a diameter of 295.4 ± 77.0 nm, and volume of 0.12 ± 0.68 μm^3^. The plasma membranes of these invaginating structures were clearly differentiated from the plasma membrane surrounding the dendritic cytoplasm (Figure 6). The size and ultrastructural appearance of these invaginated structures resembled those of dendritic and axonal filopodia^53,54^. Notably, the presence of invaginated structures within dendritic shafts is consistent with recent reports using human PFC^55^ or temporal lobe^56^ biopsy samples for VEM. This complex parent dendrite exhibited other distinctive ultrastructural features. For example, unlike nearby dendritic shafts, the dendrite with invaginated filopodia-like structures exhibited an electron-dense cytoplasm with abundant mitochondria, SER, and endosomal organelles, and received multiple Type 1 synapses (Figure 6). Altogether, these ultrastructural features indicate a pyramidal neuron dendritic region requiring greater energy production, calcium buffering and regulation, and secretory trafficking of integral membrane proteins to support enhanced synaptic neurotransmission^57^.

**Figure 5.**
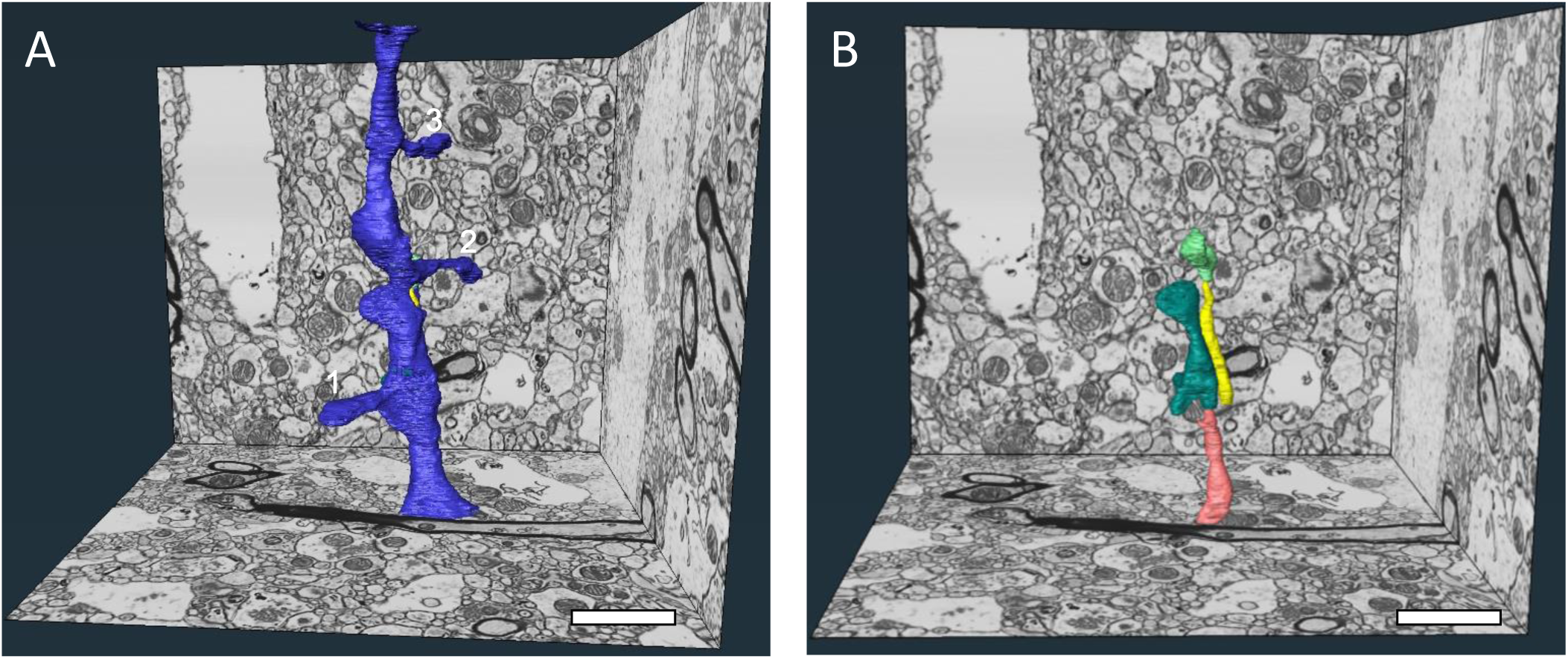
3D reconstructions of a complex dendritic shaft with filopodia-like structures coursing through the dendritic cytoplasm. **A)** A perspective overview of a volume of DLPFC layer 3 with a dendritic shaft completely reconstructed. This dendritic shaft segment spanned all 1,580 ortho-slices (7.9 μm) in the Z dimension and had a total volume of 2.0 μm^3^. Multiple spines (numbered 1-3) protrude from the parent shaft, and the neck of a fourth spine is visible at the top of the volume for a mean spine occurrence of 0.5/μm. Spines 1 and 2 each contained a spine apparatus and received a Type 1 synapse. Spine 3 also contained a spine apparatus, but was dually-innervated by a Type 1 and a Type 2 synapse. **B)** Similar perspective overview illustrating the four structures (salmon, teal, yellow and lime green) completely enveloped within the cytoplasm of the dendritic shaft. The yellow, teal and lime green axons are fully contained within the volume and are not connected to any surrounding structure. The entirety of the salmon axon is not included in the total volume. Scale bars are 2 μm.

**Figure 6.**
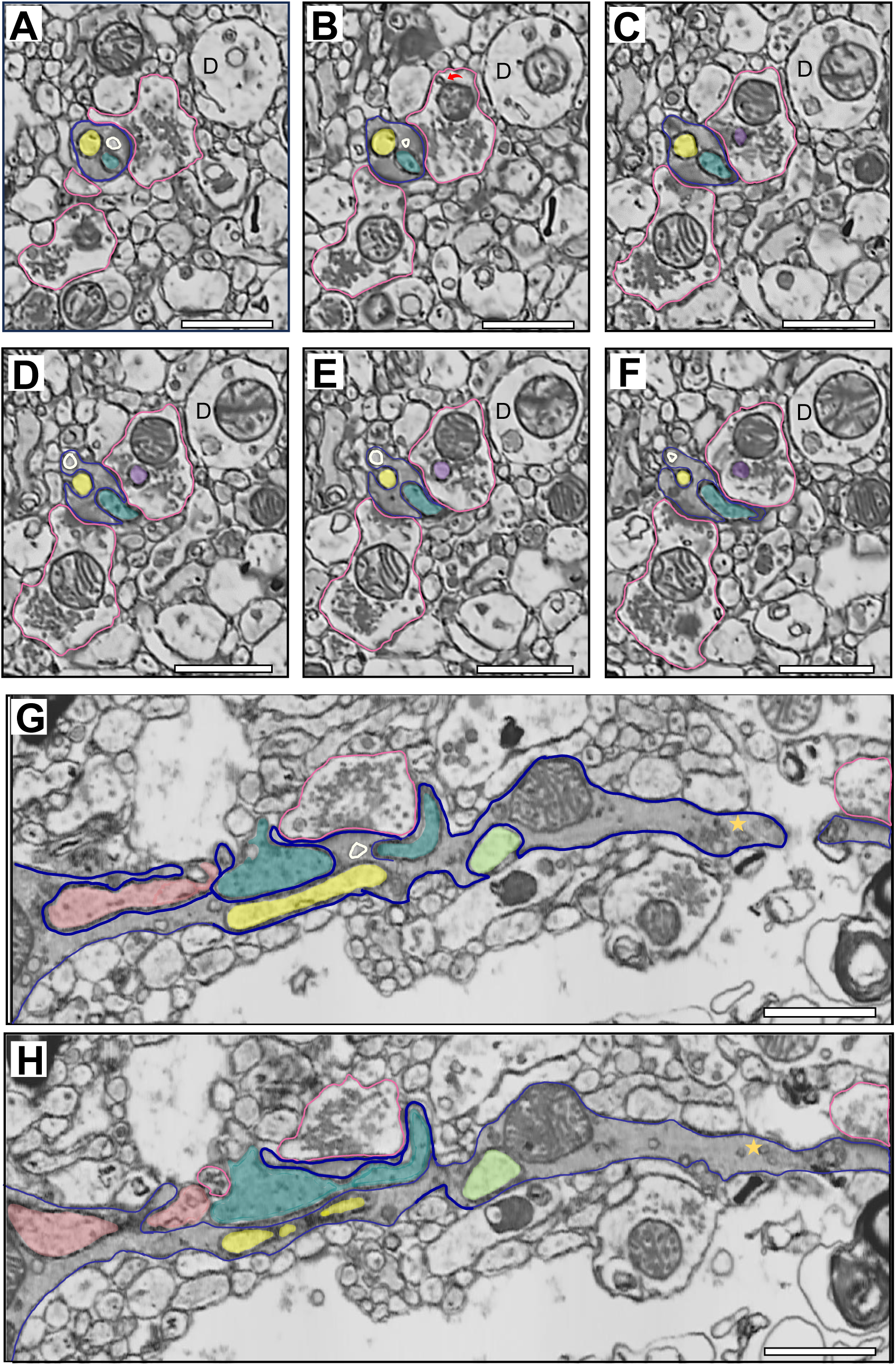
A series of electron micrographs viewed in different coordinate planes illustrating the unique ultrastructural features of the complex dendritic shaft. **A-F**) Electron micrographs spanning 315 nm in the XY coordinate plane illustrating two filopodia-like, invaginated structures (yellow and teal filled) completely enveloped within the dendritic shaft (dark blue outline) cytoplasm. The plasma membrane of the invaginated structures and the plasma membrane enclosing the dendritic cytoplasm are clearly distinct, and also show discrete, electron-dense regions which appear to be non-synaptic contacts. Unlike other dendritic profiles (**D**) in close proximity, the complex dendritic shaft exhibits an electron dense cytoplasm, indicative of an abundance of molecules involved in cellular, organelle and synaptic function. In Panel A, a single Type 1 synapse from an axonal bouton (pink outlined) is visible. Presumptive endosome large vesicle (ivory outlined) is positioned between the PSD and the two invaginated structures. In Panel B, an additional Type 1 synapse is present on the dendritic shaft. In the top bouton, a mitochondrial-derived vesicle (red curved arrow) is visible. Panels C-F illustrate the teal invaginated structure exiting the dendritic shaft, and the yellow structure partially exposed to the neuropil. Panels D-E also show an invaginated structure (purple filled) piercing the cytoplasm of the top synaptic axonal bouton. **G-H)** Electron micrographs separated by 70 nm in the XZ coordinate plane illustrating the same dendritic shaft, the Type 1 synapse initially present in Panel A, and all four filopodia-like invaginated structures (salmon, teal, yellow, and lime green filled). The electron-dense cytoplasm is apparent, as is an abundance of putative endosomal and SER structures. A classic example of amorphous vesicular clumps (gold star) is present in the dendritic shaft. Amorphous vesicular clumps likely reflect a mix of SER and endosomal vesicles^98^. The yellow and teal invaginated structures and the endosome large vesicle (ivory outlined) are visible in Panel G, positioned centrally to the PSD. D-dendrite. Scale bar is 1 μm.

### Quantitative Analysis of Densely-Reconstructed DLPFC Layer 3 Neuropil Sub-Volume

Dense reconstruction of the sub-volume (Figure 1D-E) revealed the respective contributions of each type of neuronal and glial process (Table 1). Synaptic analysis showed greater density of Type 1 synapses compared to Type 2 synapses, with an overall synaptic density of 0.39/μm^3^. Axonal boutons forming Type 1 synapses preferentially targeted dendritic spines over shafts (Table 2). Of the 28 total axonal boutons identified in the sub-volume, 71% (n=20) formed a Type 1 synapse, 18% (n=5) formed a Type 2 synapse, and 11% (n=3) did not form a synapse. To further interrogate synaptic features of postmortem human DLPFC, we completed a targeted analysis of the predominant synaptic population, Type 1 glutamatergic axo-spinous synapses.

**Table 1.** Proportion of neuronal and glial compartments present in the densely-reconstructed DLPFC layer 3 neuropil sub-volume.

| Cellular Compartment | Volume ($\mu\text{m}^3$) | % of total volume |
| --- | --- | --- |
| Unmyelinated axons | 19.1 | 29.7% |
| Dendritic shafts | 15.6 | 24.3% |
| Astrocyte processes | 9.3 | 14.4% |
| Axonal boutons | 8.2 | 12.7% |
| Myelinated axons | 3.0 | 4.7% |
| Dendritic spines | 1.4 | 2.2% |
| Extracellular space | 7.8 | 12.0% |

**Table 2.** Synaptic bouton identity, density and postsynaptic target present in the densely-reconstructed DLPFC layer 3 neuropil sub-volume.

| Bouton Type | N | Density | Postsynaptic Target |  |  |
| --- | --- | --- | --- | --- | --- |
|  |  |  | Spine | Shaft | Axon |
| Synaptic- Type 1 | 20 | 0.31/$\mu\text{m}^3$ | 14 (70%) | 6 (30%) | 0 |
| Synaptic- Type 2 | 5 | 0.08/$\mu\text{m}^3$ | 0 | 4 (80%) | 1 (20%) |
| Non-synaptic | 3 | 0.05/$\mu\text{m}^3$ | -- | -- | -- |

### Targeted Quantitative Analysis of Type 1 Synaptic, Sub-Synaptic and Organelle Structures

A total of 50 Type 1 glutamatergic axo-spinous synapses were identified and reconstructed in 3D. Quantitative analysis (Figure 7) revealed the average sizes of synaptic axonal boutons (0.46 ± 0.26 μm^3^), synaptic spines (0.15 ± 0.13 μm^3^), presynaptic active zones (5.5 ± 3.8e^6^ nm^3^), and PSDs (8.0 ± 5.1e^6^ nm^3^). Mitochondria, identified in 52% (n=26) of Type 1 synaptic boutons, had an average volume of 0.09 ± 0.05 μm^3^. All bouton mitochondria exhibited globular morphology (mean aspect ratio = 0.99). Of the synaptic spines, 88% (n=44) contained a spine apparatus. Because these core synaptic components were sufficiently preserved for quantitative volumetric and morphologic analysis, we next sought to determine whether the within– and trans-synaptic structure relationships that reflect functional synaptic communication were maintained postmortem.

**Figure 7.**
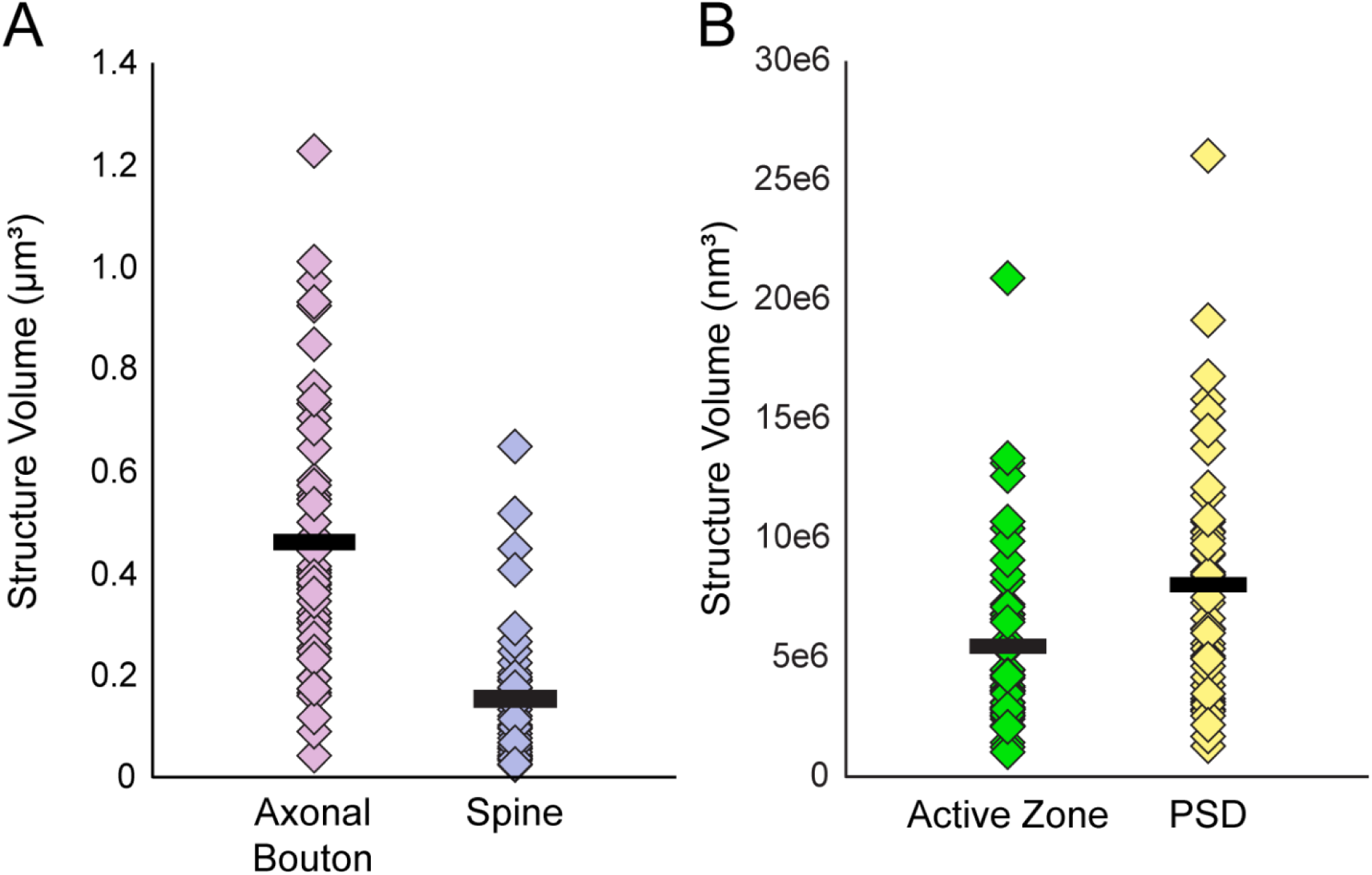
Scatter plots showing the volumes of each reconstructed synaptic and sub-synaptic structure in postmortem human DLPFC layer 3. **A)** Volumes of 50 Type 1 synaptic axonal boutons and the 50 postsynaptic spines. **B)** Volumes of the presynaptic active zones and PSDs. Markers represent an individual structure, and black bars represent the mean volume.

### Targeted Quantitative Analysis of Type 1 Pre– and Postsynaptic Structure Relationships

Presynaptic bouton and active zone volumes were positively correlated (r=0.60, p=4.0e^-6^; Figure 8A), as were spine and PSD volumes (r=0.43, p=0.002; Figure 8B). Analyses of trans-synaptic structures revealed positive correlations between presynaptic bouton and postsynaptic spine volumes (r=0.37, p=0.008; Figure 8C) and presynaptic active zone and PSD volumes (r=0.76, p=1.2e^-10^; Figure 8D). To further evaluate whether trans-synaptic ultrastructural relationships persist in the postmortem human brain, we compared synapses based on the presence of mitochondria or a spine apparatus, two organelles indicative of greater synaptic function and activity^28,29,58,59^. Type 1 axo-spinous synapses with presynaptic mitochondria had significantly greater bouton, active zone, and PSD volumes, and were significantly more likely to target a spine with a spine apparatus (Table 3). Type 1 axo-spinous synapses with a postsynaptic spine apparatus had significantly greater mean spine, active zone, and PSD volumes (Table 4). These convergent results support our assertion that the fundamental ultrastructural features reflecting synaptic function and activity previously observed in brains of model organisms are similarly preserved in postmortem human DLPFC.

**Figure 8.**
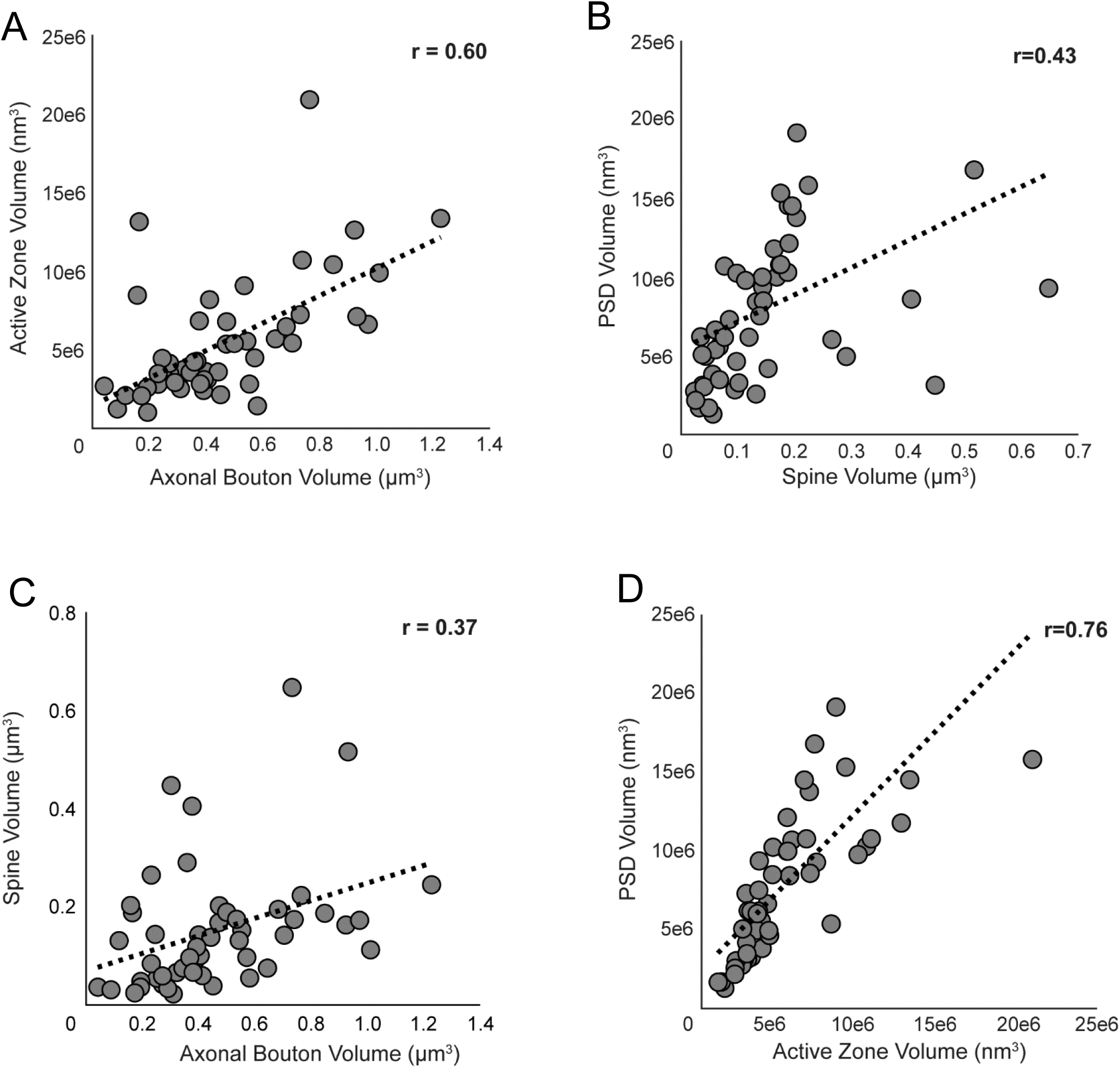
Correlation plots showing the relationships between Type 1 synaptic and sub-synaptic structures in postmortem human DLPFC layer 3. **A-D**) Markers represent the volumes of each structure within an individual synapse. Strong, positive correlations were identified between bouton and active zone volumes **(A)** and active zone and PSD volumes **(D)**. Moderate, positive correlations were identified between spine and PSD volumes **(B)** and bouton and spine volumes **(C)**.

**Table 3.**
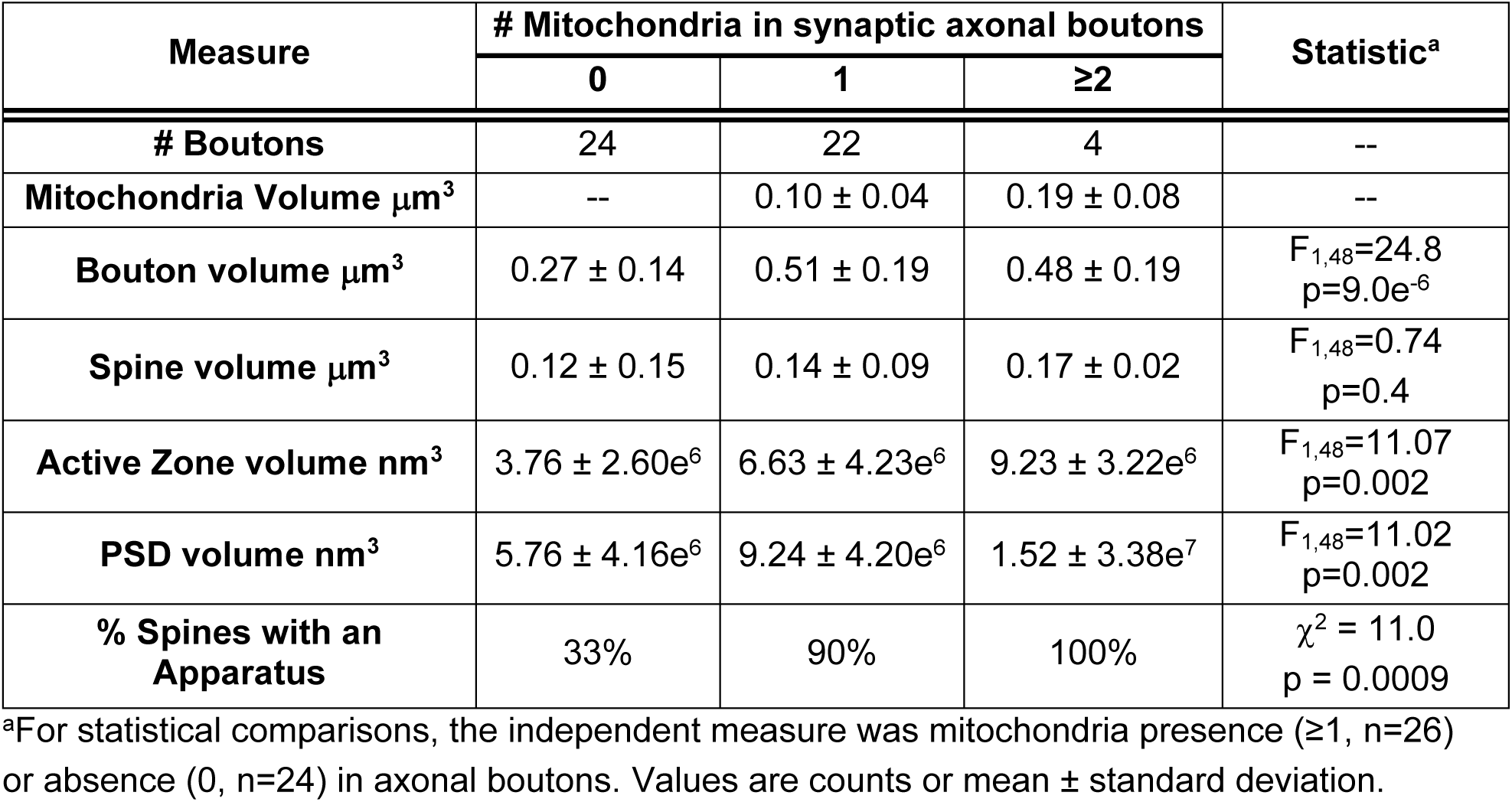
Comparison of synaptic, sub-synaptic and mitochondrial volumes based on mitochondrion abundance in Type 1 axo-spinous synapses.

**Table 4.** Comparison of synaptic, sub-synaptic and mitochondrial volumes in Type 1 axo-spinous synapses based on presence of a spine apparatus.

| Measure | Spine Apparatus |  | Statistic |
| --- | --- | --- | --- |
|  | Yes | No |  |
| **# Spines** | 44 | 6 | -- |
| **Bouton volume $\mu\text{m}^3$** | $0.41 \pm 0.21$ | $0.24 \pm 0.11$ | $F_{1,48}=3.72$, $p=0.06$ |
| **Spine volume $\mu\text{m}^3$** | $0.15 \pm 0.12$ | $0.04 \pm 0.03$ | $F_{1,48}=4.47$, $p=0.04$ |
| **Active Zone volume $\text{nm}^3$** | $5.91 \pm 3.87\text{e}^6$ | $2.25 \pm 0.89\text{e}^6$ | $F_{1,48}=5.22$, $p=0.03$ |
| **PSD volume $\text{nm}^3$** | $8.61 \pm 5.14\text{e}^6$ | $3.45 \pm 1.84\text{e}^6$ | $F_{1,48}=6.01$, $p=0.02$ |
| **% Boutons containing mitochondria** | 57% | 17% | $\chi^2 = 3.4$<br>$p = 0.06$ |
| **Mitochondria volume $\mu\text{m}^3$** | $0.11 \pm 0.06$ | 0.10 | $F_{1,48}=0.005$, $p=0.9$ |

Values are counts or mean ± standard deviation.

## DISCUSSION

We employed an advanced VEM imaging approach, FIB-SEM, to determine if 3D ultrastructural correlates of *in vivo* glutamate synapse function and activity are preserved *ex vivo* in the postmortem human brain. To address this, we first modified an existing workflow^60^ to generate postmortem human DLPFC tissue samples with optimal ultrastructural preservation and enhanced-contrast staining required for FIB-SEM imaging. Second, we successfully imaged a volume (2,630 μm^3^) of human DLPFC layer 3 with a 5 nm isotropic voxel resolution via FIB-SEM, ensuring little to no loss of information and equivalent resolution in all dimension planes. Third, we implemented a semi-automated data collection approach to complete a dense 3D reconstruction of a sub-volume of postmortem human DLPFC layer 3. Finally, we used this semi-automated approach to reconstruct sub-cellular, sub-synaptic and mitochondrial components within 50 glutamate axo-spinous synapses. Quantitative analysis of these 3D datasets revealed that ultrastructural features of synaptic function and activity were preserved within and across individual synapses in postmortem human brain tissue. Thus, our findings provide proof-of-concept that *in vivo* functioning of individual synapses in human brain can be investigated *ex vivo* in tissue obtained postmortem and analyzed by FIB-SEM.

DLPFC layer 3 neuropil organization was revealed by dense reconstruction and quantitative analysis. The relative proportion of DLPFC layer 3 neuropil occupied by each type of cellular compartment was remarkably similar to recent VEM findings in samples of temporal cortex layer 3 that were obtained by biopsy^61^, as was the volumetric proportion reflecting extracellular space in biopsy-obtained prefrontal cortex^55^. The mean synaptic density observed by VEM imaging of biopsy cortical samples^56,61–65^ ranges from 0.0002/μm^3^ to 0.74/μm^3^, and of autopsy cortical samples^62,66–71^ from 0.40/μm^3^ to 0.76/μm^3^. Our current finding of 0.39/μm^3^ in DLPFC layer 3 is consistent with these existing data, which span multiple cortical areas and layers, and is also consistent with previous findings in monkey DLPFC layer 3^72^.

We identified that 80% of DLPFC layer 3 synapses were Type 1 and the remaining 20% were Type 2. These proportions are highly consistent with findings in layer 3 of biopsied temporal cortex^61^, but lower than the findings of 92-95% Type 1 synapses in postmortem temporal cortex layers 2-5^62,67–71^, anterior cingulate cortex layer 3^67^, and layer 3 of primary visual, motor and somatosensory cortices^66^. Findings regarding the effects of PMI on Type 1 and 2 synapse identification are highly mixed [for review see^73^]. Some evidence suggests that PSD size may increase with PMI^74^, resulting in the possible mis-identification of Type 2 synapses as Type 1. However, our findings indicate this potential confound was likely not present, and/or the rich, isotropic, 3D ultrastructural data available for every synaptic contact mitigated any potential PMI effect on synapse identification. Finally, our finding that 70% of Type 1 synapses were formed onto dendritic spines is consistent with previous VEM studies of biopsy^56,64,65^ and autopsy^62,66–69^ cortical samples which found, respectively, that 82-90% and 55-73% of Type 1 synapses were formed onto dendritic spines. Thus, all neuropil components, including Type 1 and Type 2 synapses, were identifiable and quantifiable via FIB-SEM imaging using the tissue processing workflow optimized for postmortem human brain.

Targeted 3D ultrastructural analysis of Type 1 axo-spinous glutamatergic synapses similarly demonstrated that synaptic nanoarchitecture is preserved in postmortem human DLPFC. For example, in model systems, active zone and PSD sizes are highly correlated, and this finding reflects that pre– and postsynaptic components must act together as a functional unit for effective synaptic communication^34,35^. Active zone and PSD sizes are also diverse in the normal brain, which reflects the dynamic range of synaptic activity and plasticity contributing to complex neural computation^75–77^. Both of these core synaptic features were clearly evident in postmortem human DLPFC, qualitatively and quantitatively, suggesting that the synaptic functional unit remained intact postmortem.

Analysis of mitochondria and spine apparatuses, synaptic organelles whose presence is indicative of greater activity and neurotransmission^29,59^, further demonstrated preserved glutamatergic synaptic relationships within postmortem DLPFC. For example, mitochondria are trafficked to more active synaptic boutons, providing the required energetic support for efficient neurotransmitter release^78,79^. Similarly, spine apparatuses are present in dendritic spines that are more synaptically active, enhancing postsynaptic Ca^2+^ buffering and post-translational modification capabilities^58,59^. Consistent with these findings from model organisms, and a study using serial section EM analysis of postmortem human hippocampus^33^, Type 1 axo-spinous synapses with presynaptic mitochondria had significantly greater mean active zone and PSD volumes, and significantly more of these synapses contained a spine apparatus. Likewise, glutamatergic synapses with a postsynaptic spine apparatus exhibited significantly greater mean active zone and PSD volumes, and significantly more of these synapses contained presynaptic mitochondria. Together, the synaptic nanoarchitecture and organelle findings provide convergent support that relative *in vivo* synaptic function and activity can be interrogated *ex vivo* in postmortem human brain tissue via quantitative FIB-SEM imaging and analysis.

In the DLPFC layer 3 neuropil volume, we reconstructed a spiny dendritic shaft that spanned all 1,580 ortho-slices. Notably, this shaft exhibited unique and complex ultrastructural features characteristic of heightened synaptic communication, integration, and plasticity. Unlike nearby shafts, which exhibited ultrastructure features typical of dendrites in the cortical neuropil (e.g., relatively electron-lucent cytoplasm)^80^, this dendritic segment possessed an abundance of mitochondria and single-membrane organelles, such as SER and endosomes, and an electron-dense cytoplasm. Furthermore, we also identified four discrete, invaginated sub-cellular processes situated within this dendrite’s cytoplasm. Unexpectedly, these invaginating processes were not connected to any parent structure, instead presenting as independent entities within the dendritic shaft. Such a relationship suggests these processes, which resemble dendritic or axonal filopodia^53^, were trans-endocytosed by the parent dendrite^54^. Invaginating structures are implicated in synaptic formation, maintenance, pruning and plasticity, and provide a unique means to further enhance the functional flexibility of the invaginated structure^53,54,81–84^. Thus, this dendritic segment appears to be a hub of substantial postsynaptic resources that enhance the capacity for localized synaptic integration and computation^85,86^. A dendritic segment exhibiting these features has not been previously reported in either human or non-human cortex to our knowledge. The PFC is a region uniquely expanded in humans^87^, and exhibits molecular and cytoarchitectonic features thought to underlie many higher-order functions specific to humans^88^. Indeed, the uniqueness of human PFC may contribute to the apparent human-specificity of serious brain disorders^89^ such as schizophrenia^43^, autism spectrum disorder^42^ and Alzheimer’s disease^90^. Future comparative VEM studies of DLPFC neuropil will provide valuable insight into the exceptional nature of human brain.

In sum, the current findings provide proof-of-concept evidence that *in vivo* functioning of individual synapses in human brain can be investigated *ex vivo* in tissue obtained postmortem and analyzed by VEM. Our tissue processing workflow generated 3D datasets of excellent fixation, staining and ultrastructural preservation using samples with a PMI of 6.0 hours and that were in storage for more than 8 years before staining and FIB-SEM imaging. This PMI is ≥ 2 hours longer than other published VEM studies^62,66,67^, supporting that the workflow described here is compatible with longer PMIs. The success of our approach is also consistent with existing EM data demonstrating that ultrastructure is preserved, and light microscopic data of synaptic appositions, in postmortem human samples of longer PMIs [for examples, see^46,48,91–93^], and that PMI may not be the strongest predictor of ultrastructural preservation^48^. The suitability of both biopsy and autopsy human brain tissue samples, and of archived tissue samples, for quantitative VEM greatly expands the potential resource pool and opportunities to investigate ultrastructural correlates of neural function in human health and disease. The current findings show that not only are these sub-synaptic structures able to be identified, segmented, and reconstructed in 3D in postmortem human brain, but that the biological processes occurring postmortem do not affect fundamental pre– and postsynaptic relationships. Finally, our use of FIB-SEM technology has revealed a novel hub-like dendritic organization in human DLPFC. Together, these findings indicate that VEM imaging studies of postmortem human brain can be used to investigate the nature of synaptic dysfunction in brain disorders.

## LIMITATIONS OF THE STUDY

The current study is associated with some interpretative limitations. First, our analyses were completed in one subject, raising the possibility that results may not be generalizable to other subjects. Although the consistency of our results with existing EM studies in experimental model systems and previously published human brain tissue samples obtained via biopsy or postmortem (see Introduction and Discussion) suggests this likely is not the case, further studies with larger sample sizes are required. Second, quantitative VEM approaches provide an indirect way to study synaptic, sub-synaptic and mitochondrial function since direct measures of the activity and functioning of individual synapses are unavailable. Indeed, neural communication is a dynamic process, and all microscopy studies of preserved brain tissue capture only a snapshot of these processes. Relatedly, technologies with the resolving power to study the functioning of individual synapses in the *in vivo* human brain are not yet available. Future work in model systems that can first visualize dynamic synaptic processes in living brain tissue, followed by fixation and VEM imaging of the same anatomic region can more definitively correlate synaptic ultrastructure-function relationships. Third, we used a semi-automated segmentation approach that required some manual interaction. Future studies utilizing interactive deep-learning algorithms will accelerate dense segmentation and reconstruction of imaged volumes^61,94,95^. Finally, as with all studies of postmortem human brain tissue^96,97^, mitigating the effect of potential confounding factors is a key aspect of rigorous experimental design when interrogating the disease effects. In particular, our study highlights the importance of ensuring that objective measures of ultrastructural preservation and quality^48^ do not differ between comparison and disease groups.

## AUTHOR CONTRIBUTIONS

Conceptualization, J.R.G.; Methodology, J.R.G., C.B.M, K.W., and Z.F.; Formal Analysis, J.R.G.; Investigation, J.R.G., C.B.M., M.M., T.B.T., K.W., J.N., D.M.; Resources, J.R.G., D.A.L., Z.F.; Data Curation, J.R.G., M.M., T.B.T., D.M.; Writing-Original Draft, J.R.G.; Writing-Review & Editing, J.R.G., D.A.L., Z.F.; Visualization, J.R.G., C.B.M., M.M., K.W.; Supervision, J.R.G., D.A.L., Z.F.; Funding Acquisition, J.R.G., Z.F.

## ACKNOWLEDGMENTS

The authors gratefully acknowledge the digital graphics expertise of Mary Brady. Brain tissue was provided by the University of Pittsburgh Brain Tissue Donation Program. Funding support was provided by NIH grants R01MH132544 (JRG), S10MH133575 (JRG, ZF), R01ES034037 (ZF), R21AA028800 (ZF), Commonwealth of Pennsylvania Formula Fund (ZF), The Pittsburgh Foundation (ZF), and Baszucki Brain Research Funds (ZF). All data were previously published in preprint form on biorxiv.com.

## METHODS AND MATERIALS

### Human Subject

The brain specimen was obtained during a routine autopsy conducted at the Allegheny County Office of the Medical Examiner (Pittsburgh, PA) after consent for donation was obtained from the next-of-kin. An independent committee of experienced research clinicians confirmed the absence of any lifetime psychiatric or neurologic diagnoses for the decedent based on medical records, neuropathology examinations, toxicology reports, and structured diagnostic interviews conducted with family members of the decedent^99,100^. This subject was a 62-year-old male who died suddenly and out-of-hospital with an accidental manner of death. The postmortem interval (PMI, defined as the time elapsed between death and brain tissue preservation) was 6.0 hours. PFC pH was measured as 7.0, and RNA Integrity Number measured as 8.7. These demographic and tissue features are within the range of our previously published light and EM studies of postmortem human brain tissue^46,48,49,91^. All procedures were approved by the University of Pittsburgh’s Committee for the Oversight of Research and Clinical Training Involving Decedents and the Institutional Review Board for Biomedical Research.

### Tissue Preparation and Sample Selection

A tissue block approximately 1 cm^3^, containing cortical layers 1-6 and the underlying white matter, was dissected from the fresh middle frontal gyrus, Brodmann Area 46 (Figure 9A-B). The tissue block was immersed in 4% paraformaldehyde/0.2% glutaraldehyde (in 0.1M PB) for 24 hours at room temperature, followed by 24 hours at 4°C in the fixative solution. The fixed tissue block was sectioned with a 50 μm step-size on a vibratome (VT 1000P, Leica, Wetzlar, Germany), and sections were stored in a cryoprotectant solution (30% ethylene glycol/30% glycerol) at –30°C until time for histological processing.

**Figure 9.**
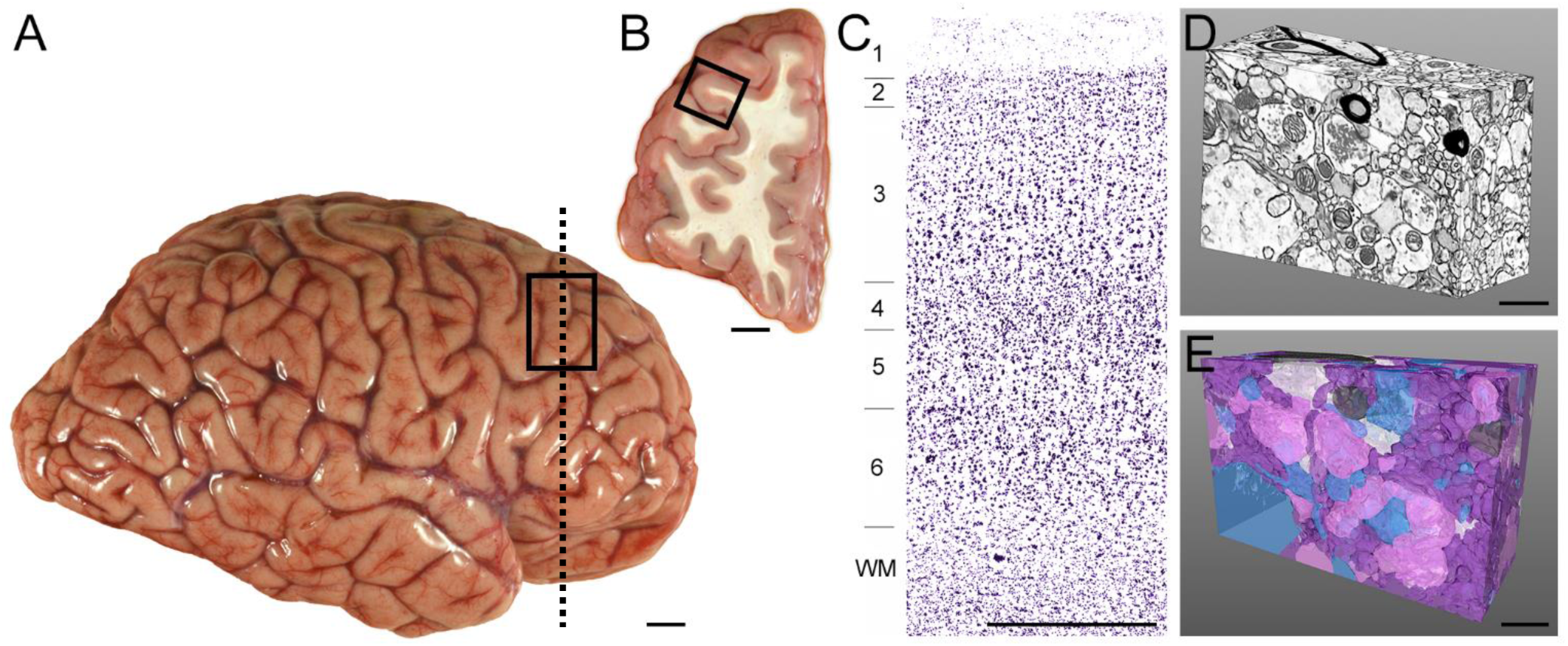
Overview of postmortem human brain tissue sampling and 3D reconstruction. **A)** Lateral view of a representative postmortem human brain hemisphere before fixation. Rectangle shows the approximate location of the DLPFC region. **B)** Coronal view of the DLPFC at the approximate rostro-caudal level indicated by the vertical dashed line in Panel A. Box indicates the approximate region sampled for this study. **C)** Nissl-stained tissue section from the approximate location shown by the box in B. Cortical layers are numbered, and tissue was sampled from layer 3 for the current study. **D)** Postmortem human DLPFC layer 3 neuropil volume imaged in 3D via FIB-SEM and analyzed in the current study. **E)** Corresponding dense 3D reconstruction of neuronal and glial profiles shown in Panel D. Glutamate axonal boutons (pink), dendritic shafts and spines (blue), un-myelinated axons (purple), myelinated axons (dark grey), and astrocytic processes (white). Scale bar in A is 2 cm, B is 1 cm, C is 1 mm, and D and E are 1 μm.

Tissue sections underwent histological processing using an approach developed by Hua and colleagues^60^ that was modified to optimize preservation, staining and contrast for postmortem human brain tissue sections. First, as briefly described above, our approach for tissue extraction, fixation, sampling, and long-term preservation substantially differed from the Hua et al. original protocol, which was designed for staining of larger-volume mouse cortex^60^. Specifically, the Hua et al. protocol collected brain tissue following transcardial perfusion of fixative buffer containing 2.5% paraformaldehyde and 1.25% glutaraldehyde, before post-extraction fixation for 24h at 4°C. In contrast, the current study employed immersion fixation since transcardial perfusion is not feasible for postmortem human brain research. We also used a lower concentration of glutaraldehyde to minimize the potential for reduced tissue antigenicity for potential future immunohistochemistry studies. For brain tissue sampling, the original Hua et al. protocol used biopsy punches of 1mm diameter and 2mm in length, whereas we performed histological staining on sections cut at 50 μm thickness to ensure that staining occurred through the entire depth of the section. Finally, given the nature and parameters of postmortem human brain tissue research, donated tissue is immediately processed and preserved using approaches that ensure long-term storage will not deleteriously affect tissue integrity. Images and data presented in the current manuscript were obtained from a tissue slice that was stored in these conditions for 8.15 years.

Modifications to the dehydration and embedding steps were also introduced, including dehydration at room temperature rather than at 4°C, and employed a different resin mixture (Electron Microscopy Sciences [EMS], Hatfield, PA, USA) containing EMbed 812, Araldite GY 502, DDSA (Dodecenyl Succinic Anhydride) and BMDA (N-Benzyl-N, N-Dimethylamine) which utilizes propylene oxide, rather than acetone, as the final dehydrant^101^. After infiltration with 100% resin, tissue sections were mounted onto glass slides coated with liquid release agent (EMS) and set at 60°C for 48 hours to allow resin polymerization. These modifications preserved the benefits of the Hua et al.^60^ approach, including a staining intensity of sufficiently high contrast to visualize neuropil components via FIB-SEM and maintenance of staining intensity throughout the depth of the tissue section.

A ∼2mm x 2mm sample of cortical layer 3 (Figure 9C) was subsequently dissected and adhered to a resin capsule. Using a Leica Ultracut UCT ultramicrotome, excess resin was trimmed with a diamond knife (Histo Diamond Knife, Diatome, Switzerland) until the face of the tissue was visible.

### FIB-SEM Imaging

The tissue sample was milled and imaged using a Helios 5 CX FIB-SEM (Thermo Fisher Scientific, Waltham, MA). After mounting onto the SEM stub using silver paste, the sample was coated with 5nm Ir for enhanced conductivity. Carbon was deposited to protect the tissue sample, and front and side trenches were milled for cross section imaging. Electron beam and ion beam fiducial markers were prepared for milling and imaging alignment. Gallium ion FIB milling and scanning electron microscopic imaging were carried out at a 52° stage tilt in an automated manner using the Thermo Scientific Auto Slice and View 4.2 Software package. FIB milling conditions were 30 kV, 0.43 nA with a slice thickness set at 5 nm. The final high-resolution backscatter electron imaging was performed at 2 kV and 0.4 nA, with a 4 µs dwell time per pixel, using the In-column detector (ICD) under immersion mode. The final image dimensions were 4055 x 3278 pixels, with a 5 nm pixel resolution. A total number of 1,580 ortho-slices were acquired at 5 nm isotropic voxels to generate a complete volume of 2,630 μm^3^ with dimensions 20.3 x 16.4 x 7.9 μm. Acquired image stacks were aligned and segmented using Amira software (Thermo Fisher Scientific).

### Ultrastructural Analysis

A sub-volume of neuropil (64.2 μm^3^ with dimensions 6275 x 4095 x 2500 nm) that excluded cell bodies and vascular structures was extracted from the master volume. Every neuronal and glial sub-cellular structure in this volume was segmented and reconstructed in 3D (Figure 9D-E), an approach termed dense reconstruction^102^, using a semi-automated approach via Amira software. In brief, the plasma membrane of each cellular structure was manually segmented in the first, middle and last ortho-slices in the XY plane. The Interpolation deep-learning module was then applied to segment the remaining plasma membrane of the structure. The operator evaluated, and corrected where needed, the segmentation in all perspective planes (XY, XZ, YZ). Because FIB-SEM imaging generates micrographic datasets with isotropic voxels, resolution is maintained in all perspective planes. As such, a 3D reconstruction of every cellular structure within the sub-volume was generated with 5nm isotropic voxel resolution. Each segmented cellular structure was evaluated in 3D and annotated as neuronal dendritic shafts, dendritic spines, axonal boutons, unmyelinated axons, myelinated axons, or glial processes using well-established ultrastructural criteria^80,103–106^.

Neuronal synaptic complexes were also identified using well-established ultrastructural criteria^80,103–106^, and Type 1 and Type 2 synaptic densities were calculated in the densely reconstructed sub-volume (64.2 μm^3^). Type 1 glutamatergic synapses were defined by a presynaptic axonal bouton directly apposed to a postsynaptic element. The postsynaptic element possessed a PSD, the electron-dense region characteristic of glutamate synapses that contains the postsynaptic proteins required for glutamatergic synaptic signaling^57^. The presynaptic bouton possessed a presynaptic active zone, defined as the electron-dense and synaptically-engaged region of the axonal bouton^107^. The pre– and postsynaptic compartments were separated by a distinct synaptic cleft^103^. Type 2, non-glutamatergic synapses were identified as described above, except the postsynaptic element did not contain a PSD, but instead exhibited greater plasma membrane electron density in the area apposed to the bouton relative to surrounding membrane. Synapses were first identified and annotated in the XY plane, then the XZ and YZ planes were evaluated to ensure that synaptic complexes were included in the synaptic density calculations, irrespective of the angle of the cut.

A targeted analysis of 50 Type 1 axo-spinous synapses was separately completed to obtain detailed volumetric measurements of glutamatergic synaptic and sub-synaptic components. Because the entirety of the synaptic complex was required to be present in X, Y, and Z dimensions for this analysis, the master volume (2,630 μm^3^) was evaluated for Type 1 synapses. Using a random Z-plane start, 50 Type 1 axo-spinous synapses were identified in the XY plane using the above-described criteria. The axonal bouton, spine head, presynaptic active zone and PSD were fully reconstructed in 3D to obtain individual ultrastructural volumes.

Each Type 1 axo-spinous synapse was evaluated for the presence of presynaptic mitochondria in boutons. Mitochondria were defined as discrete organelles identified by well-established ultrastructural criteria^80,108^, including the presence of a double membrane (comprised of mitochondrial inner and outer membranes), internal cristae membranes, and a matrix. Mitochondrial abundance^25,27,28^, size^27,29^ and morphology^30–33^ inform the relative level of activity at individual synapses^78,109^. The number of presynaptic mitochondria was tallied, and the volume of each was obtained via segmentation and 3D reconstruction. All mitochondria were classified as exhibiting globular, elongated, toroid (*i.e.*, doughnut-shaped) or damaged morphology^30,48,52^. Toroid and damaged morphologies are associated with mitochondrial dysfunction *per se* ^110,111^, and we did not observe any presynaptic mitochondria with these morphological characteristics. Globular mitochondria were identified by their spherical shape, and elongated mitochondria by a “capsule” shape^30,48,52^. The aspect ratio, defined as mitochondrion length divided by width^30^, was also calculated to confirm the morphological categorization. The aspect ratio of globular mitochondria is smaller than elongated mitochondria, and is often a value close to 1.0, given the characteristic spherical shape of globular mitochondria. For each presynaptic mitochondrion, the XY plane with the largest surface area was identified, and the length and width of the mitochondrion (measured from outer membrane to outer membrane) was measured in triplicate. The mean length measurement divided by the mean width measurement generated the aspect ratio.

Each Type 1 axo-spinous synapse was also evaluated for the presence of a spine apparatus in the postsynaptic dendritic spines. Spine apparatuses represent an extension of smooth endoplasmic reticulum (SER) into the spine head and appear as discrete tubular or vesicular structures^112,113^. Annotation as a spine apparatus required a minimum of two cisternae with an electron-dense “plate” separating cisternae in at least one ortho-slice when the spine head was evaluated in 3D^113,114^.

### Statistics

All statistical analyses were completed using SPSS software (version 27, IBM, Armonk, NY, USA). Pearson correlation coefficient analyses assessed linear relationships between structure volumes. Chi squared test determined whether the presence of mitochondria or a spine apparatus differed significantly across synaptic populations. One-way ANOVA tested differences in synaptic compartment and organelle volumes between synaptic populations.

